# Beyond Ground Truth in K-Complex Detection: A Waveform-Based SVM Classifier and the Limits of Expert Agreement

**DOI:** 10.64898/2026.05.28.728493

**Authors:** Aylin A. Vazquez Chenlo, María Cecilia Gonzalez, Laura Gorosito, Cecilia Forcato, Rodrigo Ramele

## Abstract

**Objective:** K-complexes (KCs) are large-amplitude EEG events that represent N2 sleep stage and have been linked to sensory gating, sleep protection, and memory consolidation. Their detection remains limited by inter-rater variability in visual scoring and by the reliance of detectors on features that discard temporal information. We propose a two-stage detector that combines a rule-based candidate localization algorithm with a Support Vector Machine (SVM) classifier operating directly on the raw 2-seconds waveform, and we evaluate it against an adjudicated expert consensus of two different datasets.

**Methods:** Polysomnographic recordings from 10 healthy adults (Dataset 1) were independently annotated by two human scorers; discordant events were adjudicated by a senior expert, yielding 240 consensus KCs. The automatic classifier was evaluated using subject-level 10-fold Group K-Fold cross-validation and compared directly against the two human scorers under identical conditions. Cross-dataset generalization was further assessed on the public DREAMS database (Dataset 2) under both external and internal training criteria.

**Results:** The SVM classifier achieved the highest F1-score (79.4%) and accuracy (78.8%) among all scorers, with balanced recall (81.7%) and specificity (75.8%). Of the 58 false positives, 42 originated from events both experts had rejected yet displayed canonical KC morphology and received high classifier confidence (P(KC)*>*0.7 in 45.2% of cases). This pattern was replicated on Dataset 2.

**Conclusion:** A waveform-based classifier matches expert performance and systematically flags morphologically valid KCs that fall outside conventional visual-scoring criteria.

**Significance:** These findings question the existence of an unambiguous ground truth for KC detection and support a data-driven redefinition of the event boundary, with implications for sleep staging and memory-consolidation research.

## I. Introduction

K-complexes (KCs) are large-amplitude cortical events that occur predominantly during stage N2 of non-rapid eye movement (NREM) sleep and are considered a hallmark of this sleep stage [1]. Beyond their role in sleep staging, K-complexes have been associated with multiple physiological functions, including sensory gating, sleep protection, and the regulation of cortical excitability during NREM sleep [2]–[6]. They are thought to reflect the interaction between external sensory processing and intrinsic slow oscillatory dynamics, particularly in relation to the cortical down-state [4], [5].

Growing evidence suggests that K-complexes may be involved in sleep-dependent memory processing. Neuroimaging studies indicate that spontaneous K-complexes engage thalamic and limbic networks associated with memory [7], while experimental work shows that acoustically evoked K-complexes, particularly when coupled with sleep spindles, can enhance declarative memory consolidation in humans [8].

Additionally, alterations in K-complex density and morphology have been associated with aging and cognitive decline, including neurodegenerative conditions, although their role as reliable biomarkers remains under active debate [9]–[12].

Despite their functional relevance, the identification of K-complexes remains challenging due to their morphological variability and their overlap with other slow-wave events. This variability has led to inconsistencies in visual scoring and limited inter-rater agreement, highlighting the need for reliable and scalable automatic detection methods [13], [14].

A major limitation in addressing this problem lies in the continued reliance on manual expert annotations, which is time consuming, labor intensive and difficult to scale [15]. In addition, ambiguities in morphological definitions and the overlap with other slow-wave events further compromise inter-rater reliability, challenging the notion of consistent gold standard for K-complex detection [16]–[22]. Consequently, a wide range of automatic detection algorithms has been proposed to overcome these limitations, aiming to achieve expert-level performance while improving reproducibility and scalability. These approaches typically fall into two broad categories: (1) rule-based or conventional methods [20], [23], [24], and (2) advanced signal processing techniques leveraging machine learning (ML) or neural networks (NNs) [18], [21], [25]–[28].

The first category comprises systems that rely on hand-crafted features and thresholding strategies, often using amplitude, duration, slope, and polarity characteristics. An early im-plementation included analog–digital hybrid detection schemes [23]. Matched filtering techniques have also been used to detect waveform templates in noisy signals [24], while other studies applied rule-based heuristics to define likelihood thresholds for candidate selection [20].

The second category encompasses approaches that integrate ML models or sophisticated frameworks to identify K-Complex events. These include systems based on continuous-density hidden Markov models (CD-HMMs) to model event transitions [18], classifiers trained on time–frequency representations combined with empirical rule-based selection [27], [28], graph-based clustering and one-class classification methods applied to temporal and spectral similarity patterns [26], multilayer perceptrons trained on engineered features derived from labeled K-Complex waveforms [25], or image-based classification of waveform plots using SIFT descriptors [21].

A common trend among most of these methodologies is the reliance on feature engineering, where candidate waveforms are transformed into a small set of descriptive features (e.g., peak-to-peak amplitude, duration, slope ratio) prior to classification. While this process can simplify modeling and improve generalizability, it also risks discarding subtle temporal information that may be useful for distinguishing KCs from morphologically similar slow waveforms.

In this study, we propose a two-stage approach for the automatic detection of K-Complexes, composed of a rule-based time-localization algorithm and an SVM classifier that avoids feature abstraction and instead operates directly on the full 2-second raw waveform. By preserving the complete temporal dynamics of each candidate event, the classifier can learn discriminative patterns that handcrafted features might not capture. The proposed method is evaluated against an independently adjudicated expert gold standard and directly compared with the performance of two human scorers assessed under identical conditions. Beyond detection accuracy, this work examines classification errors, revealing that a substantial proportion of the model’s false positives exhibit genuine KC-like waveform features — a finding that challenges the assumption of an unambiguous ground truth and contributes to the broader discussion on the boundaries of K-Complex definition.

## II. Methods

### A. Dataset

This study was conducted using publicly available, anonymized datasets [22], [29]. All data were originally collected with appropriate ethical approval and informed consent. No additional ethical approval was required for the present secondary analysis.

#### 1) Dataset 1

The primary dataset used for algorithm development and evaluation was derived from Experiment 2 of Forcato et al. (2020) [29], specifically from the 40-minute nap recordings of the non-reminder group. The dataset comprises polysomnographic recordings from healthy adult participants (age: 22.5± 0.7, 8 females). During the nap, participants were exposed to continuous white noise (43 dB) delivered via in-ear headphones from sleep onset until awakening. The 40-minute sleep period was defined from the first online detection of either a sleep spindle or a K-Complex visually detected by experimenter.

Polysomnographic recordings included electroencephalography (EEG), electromyography (EMG), and electrooculography (EOG) acquired using BrainAmp amplifiers (Brain Products, Munich, Germany). The EEG montage consisted of six scalp electrodes (F3, F4, C3, C4, P3, and P4, placed according to the International 10–20 System) referenced to two electrodes on the left and right mastoids. All EEG, EOG and EMG data were recorded at a sampling rate of 200 Hz and bandpass-filtered between 0.16 and 35 Hz. Polysomnographic recordings were scored offline by a trained expert according to the criteria of Rechtschaffen and Kales (1968) [30] into wake, S1, S2, S3, S4 and rapid eye movement (REM) sleep. For consistency with current nomenclature, stages were mapped to AASM criteria [1], where S1 and S2 correspond to N1 and N2, respectively, and S3 and S4 to N3.

Of the 11 available recordings, 10 were selected for this study (S23, S24, S25, S26, S27, S28, S30, S31, S32 and S33). One recording (S29) was excluded due to an insufficient number of manually detected K-complexes (N ≤ 5).

#### 2) Dataset 2

The second dataset used in this study was recorded in a hospital-based sleep laboratory in Brussels, Belgium, and is publicly available by Devuyst et al. [22]. EEG signals were acquired using a digital 32-channel polygraph (Brainnet™ System, MEDATEC) at a sampling frequency of 200 Hz. The dataset consists of 10 excerpts of approximately 30 minutes each, extracted from full-night polysomnographic recordings of healthy adult subjects. Sleep stages were scored by expert annotators according to the Rechtschaffen and Kales criteria [30], with annotations provided at 5-second resolution. Once more, for consistency with current nomenclature, stages were mapped to AASM criteria [1].

In addition to sleep stage annotations, this dataset includes expert-labeled K-complex events. K-Complexes were independently annotated by two experts. Expert 1 labeled all 10 excerpts, whereas Expert 2 annotated only the first five. All analyses were conducted using the Cz–A1 EEG derivation, and only KCs occurring during NREM stage 2 (N2) were included. In total, Expert 1 identified 212 KCs during N2 across the 10 excerpts, while Expert 2 identified 41 KCs during N2 in the five annotated excerpts (Table I).Of these, only 32 events were coincident between both experts, corresponding to an inter-rater overlap ranging from 16% to 100%.

**TABLE I.**
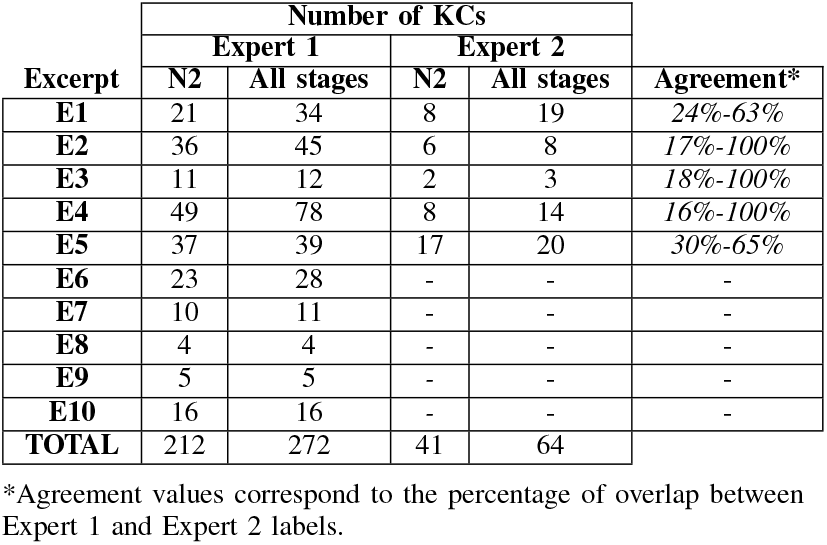
K-Complexes detected in Dataset 2 on each excerpt by each expert and their agreement*.

#### 3) Manual Annotations of K-Complex (Dataset 1)

K-Complexes were manually annotated by two expert scorers (Expert C and Expert L) under two counterbalanced labeling conditions: semi-automatic and blind labeling. In the *semi-automatic* condition, a candidate localization algorithm (Section II-C) provided initial event markers, and experts were instructed to confirm, reject, or supplement these detections. In the *blind* condition, no algorithmic guidance was provided; experts annotated both KC and non-KC events (e.g., slow waves not meeting KC criteria), maintaining an approximately balanced number of each.

To reduce scorer bias, labeling assignments were cross-counterbalanced: Expert C labeled the first five subjects in the blind condition and the remaining five using the semi-automatic approach, with Expert L following the reverse assignment. All annotations were performed on EEG channel C3, with concurrent visualization of EOG and EMG recordings, restricted to epochs scored as N2 sleep. Signal visualization has been conditioned as explained in Section II-B, consistent with standard polysomnographic scoring guidelines.

In total, Expert C annotated 175 K-Complexes and Expert L annotated 301 K-Complexes across the 10 participants (Table II). To establish a consensus ground truth, events independently identified as K-Complexes by both experts were directly accepted. For discordant annotations, a senior expert (Expert A) adjudicated the final determination. This process yielded 240 confirmed KCs across the dataset. Inter-rater agreement between Expert C and Expert L ranged from 41% to 84% depending on the subject, highlighting the substantial variability inherent in manual K-Complex scoring.

**TABLE II.** K-Complexes detected in Dataset 1 on each subject during N2 by each expert and their agreement.

|  | Number of KCs in N2 |  |  |  |
| --- | --- | --- | --- | --- |
|  | Expert C | Expert L | Final True | Agreement |
| S23 | 14 | 21 | 19 | 42% |
| S24 | 18 | 22 | 22 | 45% |
| S25 | 24 | 35 | 31 | 42% |
| S26 | 17 | 41 | 29 | 41% |
| S27 | 13 | 23 | 18 | 56% |
| S28 | 28 | 56 | 43 | 47% |
| S30 | 17 | 30 | 19 | 84% |
| S31 | 11 | 19 | 16 | 63% |
| S32 | 17 | 30 | 23 | 65% |
| S33 | 16 | 24 | 20 | 60% |
| TOTAL | 175 | 301 | 240 |  |

#### 4) Ground Truth Definitions (Dataset 2)

For consistency with Dataset 1, sleep stage annotations in Dataset 2, originally provided at 5-second resolution, were converted to 30-second epochs using a majority vote across each group of six consecutive intervals. In contrast to Dataset 1, Dataset 2 does not include a third-party adjudication to resolve disagreements between annotators. Accordingly, four labeling criteria are defined: (i) Expert 1 only; (ii) Expert 2 only; (iii) Union, where any event annotated by at least one expert is considered a K-complex; and (iv) Intersection, where only events annotated by both experts are accepted. Criteria (ii), (iii), and (iv) are restricted to the five excerpts annotated by both experts. This dataset is used both to evaluate the generalizability of the classifier trained on Dataset 1 and to independently train and assess the full detection pipeline.

### B. Pre-processing

An initial signal conditioning step was applied to preserve the key characteristics needed to distinguish K-Complexes from other EEG events while attenuating artifacts and unwanted noise. The EEG signal was filtered using a band-pass Hamming-windowed FIR filter with cut-off frequencies of 0.16 Hz and 35 Hz. The filter order was defined as 6.6 times the inverse of the shortest transition band.

The localization algorithm operates directly on the filtered EEG signal to identify potential KC candidates based on their morphological characteristics (Figure 1). The classification stage requires an additional signal conditioning step. For each candidate event, a 2-second window was extracted centered on the minimum negative point of the candidate (yellow dot in Figure 1-B and C), consisting of 1 second of preceding and 1 second of following signal. This segment was then normalized using row-wise z-score standardization: each 2-second epoch was independently scaled by subtracting its own mean and dividing by its own standard deviation (SD). This per-epoch normalization ensured that the classifier operates on the waveform shape rather than absolute amplitude, making it robust to inter-individual and inter-recording amplitude variability while preserving the temporal dynamics of each candidate event. The resulting windowed and standardized segment, comprising 400 samples at 200 Hz, served as the input feature vector to the classification model.

**Fig. 1.**
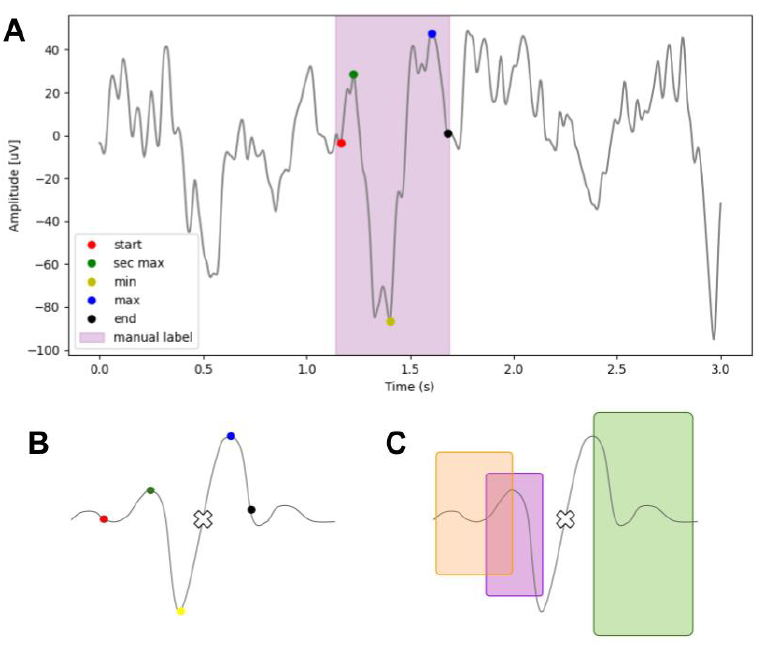
K-Complex characterization. **A** Real K-Complex manually labeled with the automatic algorithm overlay showing the detected characteristic points. **B** Ideal K-Complex waveform with feature points marked (white X indicates the center). **C** Ideal K-Complex regions of interest: orange for the start point, purple for the second maximum, and green for the end point.

### C. Time-Localization Algorithm

The algorithm used to identify candidate K-Complexes (KCs), both for semi-automatic expert labeling and as input to the subsequent machine learning classifier, is based on a sliding window approach which analyzes different characteristics. A 1-second moving window with a step size of 0.1 seconds scans the EEG signal and if no candidate is detected, the window shifts forward. The algorithm follows the steps outlined below:

1. The algorithm proceeds only if the current window corresponds to stage N2 sleep (S2). Otherwise, the window is skipped.
2. Within the window, the algorithm checks for KC-like morphology: A negative peak followed by a positive peak (i.e., minimum precedes maximum); A peak-to-peak amplitude exceeding 75*µV*; A duration between the minimum and maximum exceeding 0.5 seconds. If these criteria are not met, the candidate is discarded.
3. If the criteria are met, the algorithm proceeds to locate and define the candidate KC event using the following steps:
  a. The event is temporally centered between its minimum and maximum points, expanding to a 3-second window, ±1.5 seconds around the center (Figure 1-A).
  b. This 3-second segment is band-pass filtered using a 5th-order Butterworth filter (implemented via second-order sections), with cut-off frequencies at 1–8 Hz to preserve slow-wave and delta activity.
  c. The minimum value is identified within the 1.0–1.45 second interval of the filtered window (Figure 1-C).
  d. A second maximum is sought between the minimum and the preceding 0.3 seconds (i.e., within the 0.5–1.0 second interval) (Figure 1-B).
  e. The start of the event is defined as the nearest zero-crossing (from negative to positive) prior to the second maximum (Figure 1-C).
  f. The final maximum is located in the 1.55–1.8 second interval of the window *(Figure 1-B)*.
  g. The event’s end is defined as the nearest zero-crossing (from positive to negative) after the final maximum (Figure 1-C).
  h. The duration between start and end points is checked. If it is shorter than 0.5 seconds, the segment is symmetrically extended to meet the minimum duration. If it exceeds 2 seconds, it is trimmed symmetrically to match the upper limit.
4. If all the above points are successfully identified and validated, the segment is marked as a KC candidate and proceeds to the end of the identified event. Otherwise, the algorithm proceeds to the next window (0.1 seconds step).

### D. Classification Algorithm

#### Model selection

Seven classifiers were evaluated on Dataset 1: Support Vector Machine (SVM) with radial basis function (RBF) kernel, *ν*-SVM, Gaussian Naive Bayes, XGBoost, Random Forest, Ridge Classifier, and Logistic Regression. All models were trained on the consensus-labeled subset of Dataset 1 (240 KC and 240 non-KC samples). Model comparison was performed using Group K-Fold cross-validation with subject-level grouping to prevent data leakage between participants. SVM achieved the best trade-off between recall and precision, and was therefore selected as the final classifier. Detailed results for all seven models are provided in the Supplementary Material (Section S1).

#### Hyperparameter optimization

For the selected SVM classifier with RBF kernel, hyperparameter tuning was conducted using Stratified Group K-Fold cross-validation (5 splits) over the consensus-labeled subset (480 samples). A grid search was performed over the regularization parameter *C ∈* {0.1, 0.5, 1, 2, 5, 10, 20} and the kernel coefficient *γ ∈* {scale, 0.01, 0.005, 0.001} where 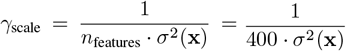, with class weighting set to balanced. The optimal configuration was selected based on the balance between recall and precision across folds, yielding *C* = 2.0 and *γ* = scale

#### Final evaluation

The optimized SVM classifier was evaluated using Group K-Fold cross-validation with 10 splits, ensuring that all samples from the same subject appeared exclusively in either the training or test set within each fold. This subject-level partitioning provides a rigorous assessment of generalization to unseen individuals. Performance was assessed using recall, specificity, accuracy, and F1-score, computed both as averages across folds (with SD) and as global metrics derived from the aggregated confusion matrix.

#### Experimental configurations

Six experimental configurations were defined: (1) *Annotations on Dataset 1 — Consensus:* The classifier trained and tested in 10-fold Group K-Fold cross-validation of the consensus-labeled subset of Dataset 1 (240 KC, 240 non-KC) with direct comparison against Expert C and Expert L. (2) *Annotations on Dataset 1 — Non-consensus:* The classifier trained in all consensus samples of Dataset 1 and predicted the 3091 non-consensus candidates to test whether the model generalizes beyond clear-cut cases to morphologically ambiguous events. (3) *Annotations on Dataset 2 — External test:* The classifier trained on Dataset 1 was applied directly to Dataset 2, under both the Expert 1 criterion and the union criterion, providing a measure of cross-dataset generalizability. (4) *Annotations on Dataset 2 — External test on Union criterion:* Same configuration as 3 with KCs defined by either Expert 1 or Expert 2. (5) *Annotations on Dataset 2 — Internal test on Expert 1 criterion:* The full pipeline (localization + classification) was trained and evaluated within Dataset 2 labeled by Expert 1 using 5-fold Group K-Fold cross-validation with a random 3:1 undersampling to mitigate class imbalance between non-KC to KC. (6) *Annotations on Dataset 2 — Internal test on Union criterion:* Same configuration as 5 with KCs defined by either Expert 1 or Expert 2.

In addition, the experimental configuration 1 (Annotations on Dataset 1 — Consensus) was also studied in four sub-analyses: events where both experts scored KC (Both KC), events where both scored non-KC (Both non-KC), events where the experts disagreed (Disagreement), and events where one or both experts did not provide a score (Not scored).

### E. Performance Evaluation Metrics

The performance of the localization and classification stages was assessed using complementary metrics derived from the confusion matrix, where TP, TN, FP, and FN denote true positives, true negatives, false positives, and false negatives, respectively. For the classification algorithm, four metrics were employed:

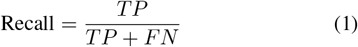

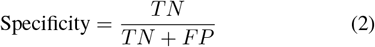

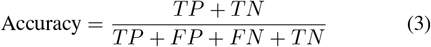

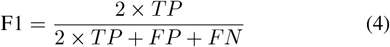

For the balanced Dataset 1, all four metrics are reported. For Dataset 2, recall and specificity are emphasized as they are independent of class prevalence. Classification metrics were computed as the mean and SD across cross-validation folds, and as global metrics derived from the aggregated confusion matrix (summing TP, TN, FP, and FN across all folds). The performance of Expert C and Expert L was evaluated against the same gold standard (Expert A) using identical metrics.

### F. Computational Setup

All analyses were implemented in Python 3.9.13 and executed on a Lenovo ThinkPad E490 equipped with an Intel Core i7 (8th generation) processor at 1.80 GHz, 8 GB RAM, and a solid-state drive. The primary libraries used were scikit-learn [31] for machine learning, MNE-Python [32] for EEG signal processing, and imbalanced-learn [33] for resampling strategies. Source code and scripts are publicly available at https://github.com/aylinavch/KCdetectionalgorithm 1001[34].

## III. Results

### A. Localization Performance

The localization algorithm successfully identified all 240 expert-annotated KCs in Dataset 1 (100% recall on the 240 consensus KCs), generating a total of 3571 candidate events (240 KC, 3331 non-KC). On Dataset 2, the algorithm detected 87% of KCs annotated by Expert 1 (184 out of 212) and 80% of those annotated by Expert 2 (33 out of 41), producing 5329 candidate events across the 10 excerpts. Table III summarizes the localization results for each experimental configuration.

**TABLE III.** Performance of the localization algorithm in each experiment configuration.

| Experiment Configuration | Excerpts/ Subjects | Annotated KC | Localized KC | Total non-KC |
| --- | --- | --- | --- | --- |
| 1 | 10 | 240 | 240 | 240 |
| 2 | 10 | 240 | 240 | 3331 |
| 3 & 5 | 10 | 212 | 184 | 5044 |
| 4 & 6 | 5 | 163 | 133 | 2415 |

### B. Classification Performance on Dataset 1

The SVM classifier was evaluated on Dataset 1 consensus-labeled dataset using 10-fold Group K-Fold cross-validation (Table IV). The classifier achieved a mean recall of 81.2% (±12.0%; global of 81.7%) and specificity of 75.8% (±8.2%; global of 75.8%), both intermediate between the two human experts. The classifier achieved the highest accuracy of 78.8% (±7.4%; global of 86.9%), and F1-score of 79.3% (±8.1%; global of 79.4%) across folds. Global metrics were computed from the aggregated confusion matrix: TP = 196, TN = 182, FP = 58 and FN = 44.

**TABLE IV.**
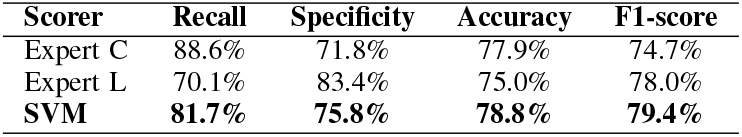
Performance of the classification algorithm in Dataset 1.

The relatively high standard deviation across folds, particularly for recall (±12.7%), reflects substantial inter-subject variability in KC morphology and detection difficulty. The majority of subjects achieved accuracy above 75%, with one notable exception: Subject S27 (accuracy = 58.3%, recall = 55.6%, specificity = 61.1%), which is described in the Supplementary Material (Section S2).

### C. Error Analysis

To better understand the relationship between misclassifications and the degree of agreement between human scorers, model’s classification accuracy and error distribution were calculated across the 4 sub-analysis of experimental configuration 1 (Table V).

**TABLE V.** Accuracy of the detection algorithm stratified by inter-rater agreement category of experiment configuration 1.

| Category | Total number of events | Accuracy | FP | FN |
| --- | --- | --- | --- | --- |
| Both KC | 126 | 88.9% | 0 | 14 |
| Both non-KC | 124 | 66.1% | 42 | 0 |
| Disagreement | 115 | 73.0% | 1 | 30 |
| Not scored | 115 | 87.0% | 15 | 0 |

The model performed best on events where both experts agreed on a KC label (88.9% accuracy). The most informative category is “Both non-KC”, where both experts independently rejected the events as KCs, yet the model classified 42 of 124 (33.9%) as positive. In the “Disagreement” category, the model missed 30 of the 115 events (26.1% FN rate), indicating that events where experts themselves disagree pose the greatest challenge for automated classification.

### D. False positive Morphology Analysis

The 42 false positives arising from the “Both non-KC” category were examined in detail. The True KC average displays the canonical KC morphology: a sharp negative deflection near 1.0 s followed by a slower positive rebound. The False positive average follows the True KC waveform shape, exhibiting a negative-positive biphasic pattern, albeit with reduced amplitude (Figure 2). Individual False positive waveforms confirmed that this similarity is consistent across the majority of False positive events.

**Fig. 2.**
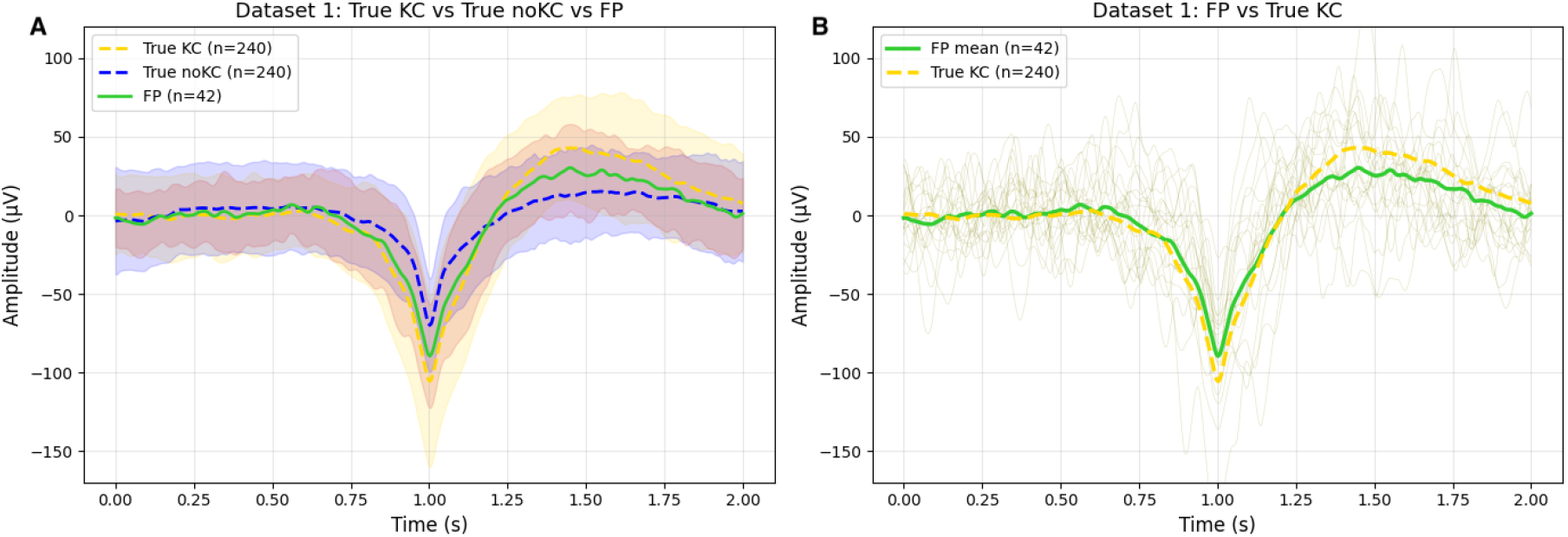
Average waveforms for true KC, true non-KC, and false positive (FP) from the “Both non-KC” category in Dataset 1. **A** Average for all the events. **B** Individual FP waveforms overlaid with FP and true KC averages. Shaded regions represent mean ±1 SD.

The posterior probability assigned by the classifier *P*(*KC*) was examined across all four outcome categories. True positives and true negatives were well separated (median *P*(*KC*) = 0.82 and 0.15, respectively). False positives exhibited a mean *P*(*KC*) of 0.69 (median = 0.68), with 45.2% receiving *P*(*KC*) *>* 0.7. False negatives showed a mean *P*(*KC*) of 0.27. These findings indicate that the classifier’s false positives are not marginal cases near the decision boundary; rather, the model assigns them moderate to high confidence as KC events (Figure 3).

**Fig. 3.**
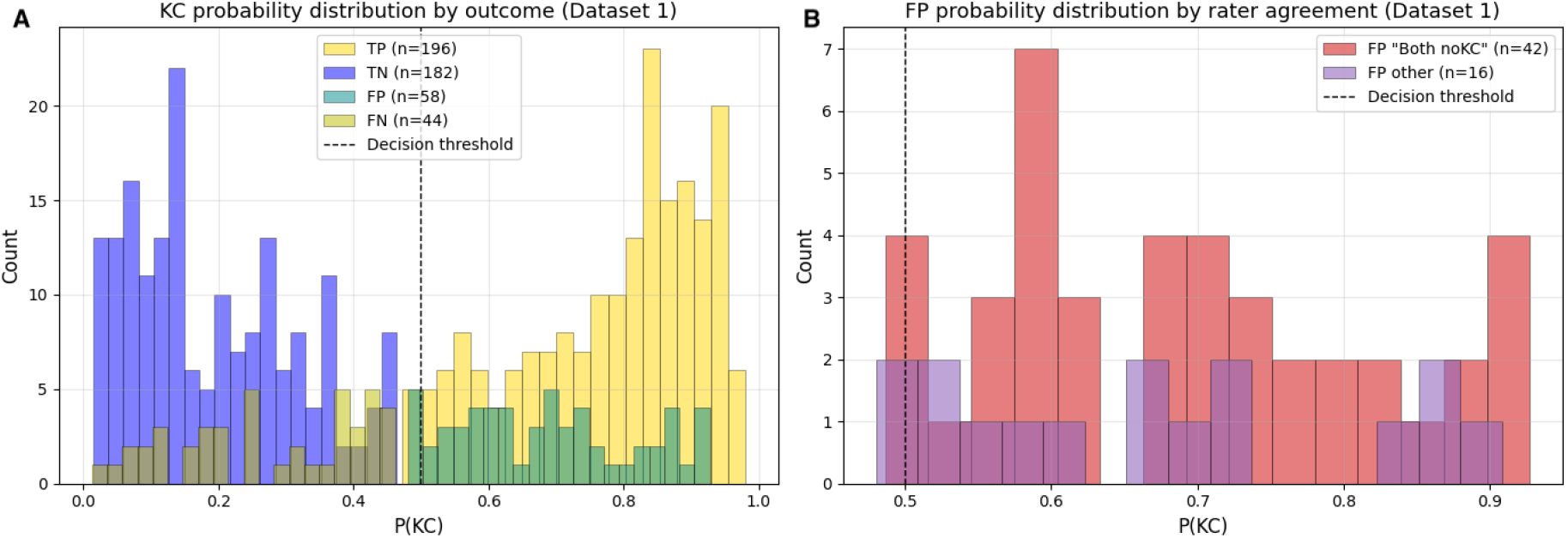
Distribution of probability for the SVM-classifier by outcome category in Dataset 1. **A** All four categories: true positive (TP), true negative (TN), false positive (FP) and false negative (FN). **B** FP events stratified by inter-rater agreement (“Both non-KC” vs. other FP). Dashed line indicates the decision threshold (0.5).

### E. Cross-Dataset Evaluation on Dataset 2

#### External evaluation

The SVM classifier trained on all 480 consensus-labeled samples from annotations on Dataset 1 was applied to Dataset 2. The model achieved a recall of 54.4% and specificity of 85.8% under the Expert 1 criterion, and a recall of 54.8% and specificity of 88.1% under the union criterion (Table VI). The substantial drop in recall relative to the primary dataset (81.7%) indicates limited cross-dataset generalization, while the preserved specificity suggests that the model’s representation of non-KC events transfers more robustly. The same morphological analysis applied to the primary dataset was performed. Figure 4 presents the average waveforms of false positives and false negatives alongside Dataset 1 and Dataset 2 KC and non-KC averages.

**TABLE 6.**
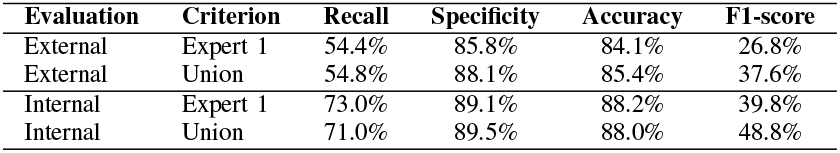
Classification performance in Dataset 2, comparing External and Internal evaluation configuration.

**Fig. 4.**
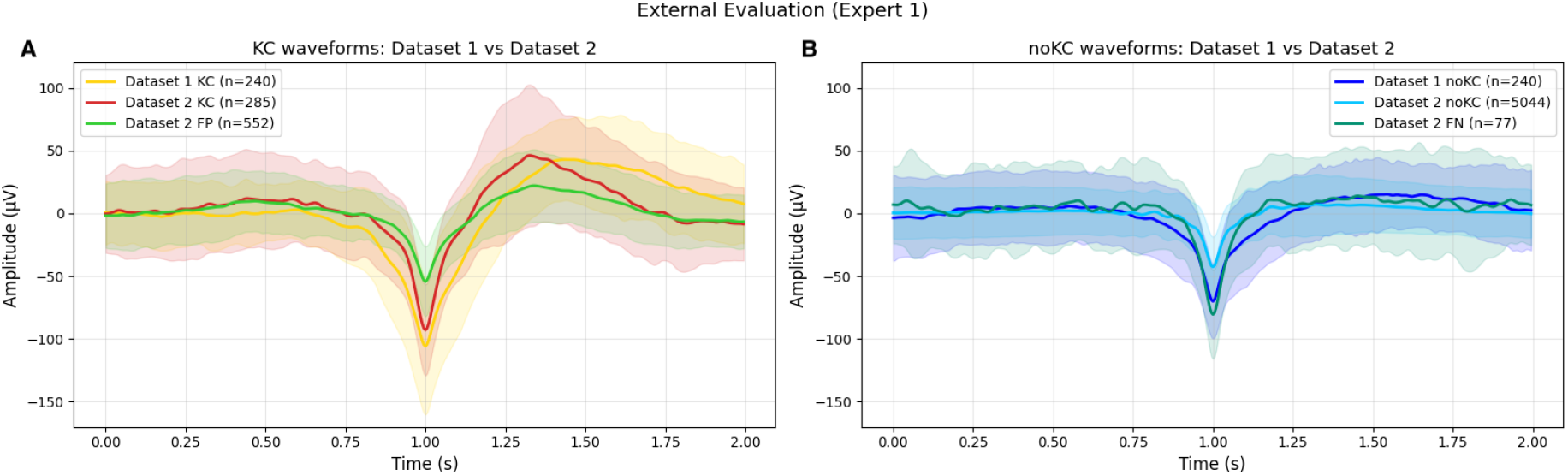
External Evaluation on waveform morphology (Expert 1 criterion). **A** FP average compared against Dataset 1 and Dataset 2 KC averages. **B** FN average compared against Dataset 1 non-KC and Dataset 2 non-KC averages. Shaded regions represent ±1 SD.

#### Internal evaluation

When the full pipeline was trained and evaluated within Dataset 2 using 5-fold Group K-Fold cross-validation with undersampling, performance improved substantially. Under the Expert 1 criterion, the model achieved a global recall of 73.0%, specificity of 89.1%, and F1-score of 39.8%. Under the union criterion, global recall reached 71.0%, specificity 89.5%, and F1-score 48.8% (Table VI). The higher F1-score under the union criterion reflects the more favorable class ratio when both experts’ annotations are pooled. The low F1-score values in both cases are attributable to the severe class imbalance in the test sets (5-8% KC prevalence), which inflates the absolute number of false positives even at high specificity, reducing precision substantially. Recall and specificity, being prevalence-independent, provide a more informative assessment under these conditions. Figure 5 presents the average waveforms of false positives and false negatives alongside Dataset 2 true KC and true non-KC averages.

**Fig. 5.**
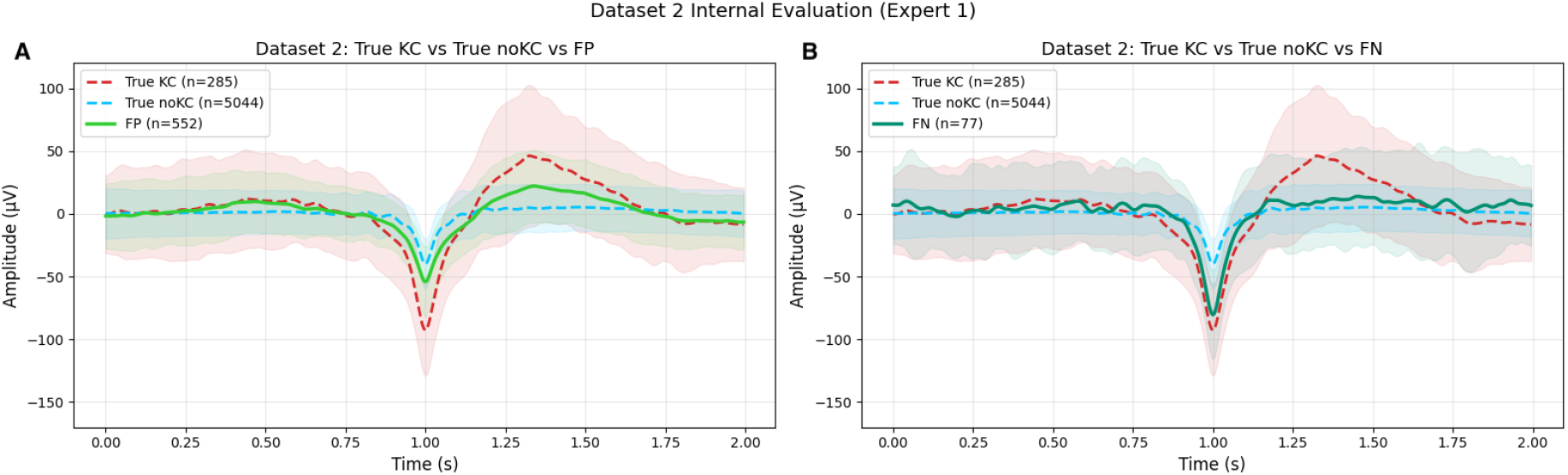
Internal Evaluation on waveform morphology (Expert 1 criterion). **A** FP average compared against true KC and true non-KC averages. **B** FN average compared against true KC and true non-KC averages. Shaded regions represent ±1 SD.

Equivalent analyses were conducted using the union labeling criterion for both external and internal evaluation, yielding morphologically consistent results. The corresponding waveform figures for the union criterion are provided in the Supplementary Material (Section S3 and S4).

## IV. Discussion

The present study demonstrates that a waveform-based classifier operating directly on raw EEG signals can achieve performance comparable to, and in some metrics exceeding, human experts, while systematically identifying morphologically valid K-complex–like events that fall outside conventional visual scoring criteria. These findings highlight both the potential of data-driven approaches for automated detection and the limitations inherent to current definitions of K-complexes based on visual inspection.

A central aspect of this work is the departure from traditional feature engineering. The majority of existing KC detection methods rely on handcrafted features extracted from candidate waveforms prior to classification. Bankman et al. [21] defined 14 amplitude- and duration-based features; Al-Salman et al. [37]–[39] extracted multi-domain features; Hernández-Pereira et al. [35] demonstrated that feature selection could improve accuracy while reducing dimensionality. While these approaches have achieved high reported accuracy, in some cases exceeding 97%, they inherently discard temporal information. In contrast, the present method preserves the full temporal structure of each waveform by using the raw 2-second EEG segment as input to the classifier. This strategy avoids potential information loss and allows the model to learn discriminative patterns directly from the data, resulting in competitive performance without the need for explicit feature design.

Importantly, direct comparison with previously reported results should be interpreted with caution (e.g., 97% accuracy [37], [39]). As previously discussed by several authors [15], [27], [35], [39], differences in evaluation protocols, class balance strategies, cross-validation schemes, and ground truth definitions make fair comparison across studies difficult. In the present work, a subject-level Group K-Fold cross-validation scheme was employed to prevent data leakage, providing a more conservative and realistic estimate of performance.

This approach is further supported by recent work highlighting the risks of excessive preprocessing in EEG analysis. Delorme (2023) [40] demonstrated that commonly used pre-processing techniques had no positive impact or may even reduce statistical sensitivity in ERP studies. By relying on minimally processed data and learning directly from the signal itself, this method avoids potential distortions while maintaining competitive or superior classification performance.

Notably, the classifier achieves performance comparable to that of human experts. A central contribution of this work is the direct comparison of the automated classifier against two independent human experts, all evaluated against the same adjudicated gold standard. The SVM classifier achieved the highest F1-score (79.4%) and accuracy (78.8%) among the three scorers. Expert C exhibited high recall but low specificity (88.6% - 71.8%) indicating a liberal scoring tendency with more false positives, while Expert L showed the opposite pattern (70.1% - 83.4%) reflecting a more conservative scoring approach. The classifier achieved a balanced trade-off between these two tendencies. The error pattern analysis revealed that no single event was misclassified by all three scorers simultaneously, suggesting that an ensemble approach combining automated and manual scoring could potentially outperform any individual scorer alone.

Of the 58 events misclassified as KC by the model, 42 (72%) originated from the “Both non-KC” category — events independently rejected by both experts. Yet these events exhibited clear KC-like morphology (*Figure 2*) and received moderate to high classifier confidence (mean *P*(*KC*) = 0.69; 45.2% with *P*(*KC*) *>* 0.7). This pattern was replicated on Dataset 2 in both external (Figure 4) and internal (Figure 5) evaluations. These findings suggest that the classifier systematically detects events with genuine KC morphological characteristics that fall below the detection threshold of visual scoring. This raises a fundamental question: are these false positives truly non-KC events, or do they represent a KC-like activity that current visual scoring criteria are too restrictive to capture?

This interpretation aligns with the broader literature on inter-rater variability in KC detection. Krohne et al. [36] reported an agreement rate of only 33% between visual scorers on Dataset 2; in our primary dataset, inter-rater agreement ranged from 41% to 84% across subjects. Such variability reflects genuine ambiguity in the boundary between K-Complexes and other slow-wave events. Automated systems that learn from the complete waveform may ultimately provide a more consistent and reproducible basis for KC identification. Several limitations should be acknowledged. First, the primary dataset is relatively small (10 subjects, 480 labeled events). Second, cross-dataset generalization was limited: the model trained on Dataset 1 achieved only 54% recall on Dataset 2, attributable to morphological differences in the candidate pools between datasets (Figure 4). Third, the classifier operates on single-channel EEG segments without access to contextual information such as sleep stage transitions or multi-channel spatial patterns. Incorporating such features — potentially through recurrent or attention-based architectures — could improve discrimination of ambiguous cases. Future work should investigate whether these false positives represent a distinct electrophysiological category or low-amplitude KCs below conventional scoring thresholds.

In summary, this work provides both a methodological and conceptual contribution. By demonstrating that waveform-based classification can match expert performance while revealing systematic limitations of visual scoring, it supports the need to move beyond rigid definitions of K-complexes. A more nuanced, data-driven characterization of these events may be crucial for clarifying their functional role within sleep architecture, particularly in relation to sensory gating, sleep protection, cortical excitability, and memory-related processes.

## V. Conclusion

This work presented a two-stage automatic K-Complex detection framework that combines a rule-based localization algorithm with an SVM classifier operating directly on raw EEG waveforms. Evaluated on Dataset 1 against an adjudicated expert gold standard, the classifier achieved the highest F1-score (79.4%) and accuracy (78.8%) among all scorers, including two independent human experts, while maintaining balanced sensitivity (81.7%) and specificity (75.8%).

Systematic error analysis revealed that the majority of the classifier’s false positives exhibit clear KC-like morphology and receive high classification confidence, suggesting that they may represent morphologically valid K-Complexes that fall outside the boundaries of conventional visual scoring. This finding, replicated on the independent Dataset 2, challenges the assumption that expert annotations constitute an unambiguous ground truth and motivates the development of more objective, data-driven criteria for KC definition.

Beyond its methodological contribution, the proposed framework offers a scalable and reproducible tool for K-Complex identification in both research and clinical contexts. Potential applications include automated scoring of large polysomnographic datasets, longitudinal monitoring of sleep architecture in aging and neurodegenerative conditions, and quantitative characterization of KCs in studies of sensory gating and sleep-dependent memory consolidation. By operating directly on the raw waveform and exposing the limits of expert agreement, this approach supports a re-examination of the operational definition of K-Complexes and provides a foundation for future investigations into their distinct physiological role within NREM sleep regulation.

## Supporting information

Supplementary Material of the manuscript

## Acknowledgment

The authors had full access to all data and code used in this study and take complete responsibility for the integrity of the analyses and the interpretation of results. The authors acknowledge the use of LLM for language editing and grammar refinement. No content was generated by AI for the technical, methodological, or analytical sections; all results, code, and scientific interpretations are the sole work of the authors.

## Notes

### Competing Interest Statement

The authors have declared no competing interest.

