## Supplementary Material of the manuscript for "Beyond Ground Truth in K-Complex Detection: A Waveform-Based SVM Classifier and the Limits of Expert Agreement"

Aylin A. Vazquez Chenlo*†, María Cecilia Gonzalez *, Laura Gorosito *, Cecilia Forcato* and Rodrigo Ramele †
* Laboratorio de Sueño y Memoria, Depto. de Ciencias de la Vida, Instituto Tecnológico de Buenos Aires (ITBA)
Buenos Aires, Argentina
† Laboratorio de Neurotrónica, Depto. de Ingeniería Informática, Instituto Tecnológico de Buenos Aires (ITBA)
Buenos Aires, Argentina

**

### S1. Model Selection: Best Machine Learning paradigm.

Six paradigms of classification were evaluated to determine the most suitable approach for automated K-complex (KC) detection. The selection of these paradigms was motivated by their prevalence and demonstrated efficacy in the literature [1]–[5].

Support Vector Machine (SVM) constitute the most widely evaluated paradigm in K-complex classification [2], [3], [6]. By projecting input features into a higher-dimensional space via kernel functions, SVMs construct a maximum-margin hyperplane that enables effective separation even when class boundaries are nonlinear [2]. Three kernel functions were evaluated: the Radial Basis Function (RBF), which has consistently yielded strong results across EEG classification tasks [5], [6]; the polynomial kernel, which captures interaction effects among features [2]; and the linear kernel, which provides a simpler, regularized decision boundary useful as an upper-bound baseline for linearly separable representations [3]. Both the standard SVM (SVC) and the $\nu$-SVM (NuSVC) formulations were included, the latter offering an alternative parameterization that bounds the fraction of support vectors and margin errors [2].

Logistic Regression and Ridge Classifier were included as representative linear models. Logistic Regression has been benchmarked in prior KC classification studies and provides probabilistic outputs amenable to threshold tuning [2], [5], [7]. The Ridge Classifier, which reframes binary classification as a regularized least-squares problem, has been proposed in the K-complex detection literature as a computationally efficient linear alternative [5].

Random Forest and XGBoost were evaluated as ensemble strategies. Random Forest, which aggregates predictions from multiple decision trees trained on bootstrap samples, has been applied to KC and sleep EEG classification and demonstrates robustness to overfitting [1], [5]. XGBoost, a gradient-boosted tree framework with regularization, has shown competitive performance in biomedical signal classification tasks [4], and was included to assess whether boosting-based aggregation offers advantages over bagging under the imbalanced conditions of the present dataset.

Gaussian Naïve Bayes was included as a generative probabilistic baseline. Despite its conditional independence assumption, Naïve Bayes has been shown to be an effective and efficient classifier for EEG-based KC detection [1], [6], [8], making it a relevant point of comparison against discriminative approaches.

In the other hand, two reference methods were included to anchor the performance range. K-Means clustering was employed as an unsupervised baseline, following prior work that used it as a reference classifier in EEG event detection [1], [6], [9]. Thresholded Linear Regression was included as a minimal supervised baseline, reflecting the class of simple feature-thresholding approaches historically used in K-complex detection [2], [10], [11].

All models were trained on the consensus-labeled subset (240 KC and 240 non-KC samples, corresponding to events where expert agreement was established or adjudicated by Expert A). All supervised classifiers used balanced class weights to account for the KC/non-KC class imbalance. Model comparison was performed using 5-fold Stratified Group K-Fold cross-validation with subject-level grouping to prevent data leakage between participants. Model selection was performed penalizing false negatives (missed KCs) more heavily than false positives, reflecting the asymmetric cost of failing to detect a K-complex in a clinical context [10]. Recall and precision on the KC class were therefore reported as the primary evaluation criteria, alongside a utility score combining both.

SVM with RBF kernel achieved the best trade-off between recall ($83.5 \pm 8.5$ %) and precision ($76.8 \pm 4.3$ %), and the highest mean utility across folds, and was therefore selected as the final classifier for hyperparameter optimization. This result is consistent with prior findings by [2], who reported that an SVM with RBF kernel achieved the best overall performance

TABLE I
PERFORMANCE OF THE 6 PARADIGMS CLASSIFIERS ON THE CONSENSUS-LABELED SUBSET, ASSESSED UNDER 5-FOLD STRATIFIED GROUP K-FOLD CROSS-VALIDATION WITH SUBJECT-LEVEL GROUPING.

| Paradigm & Model | Recall (KC) | Precision (KC) | FP (avg / total)* | FN (avg / total)* | TP (avg / total)* | TN (avg / total)* |
|---|---|---|---|---|---|---|
| ***Kernel-based SVMs*** | | | | | | |
| SVM (rbf) | $\mathbf{83.5 \pm 8.5}$ % | $\mathbf{76.8 \pm 4.3}$ % | 11.8 / 59 | 8.8 / 44 | 39.2 / 196 | 36.2 / 181 |
| NuSVC (rbf) | $80.5 \pm 7.8$ % | $75.0 \pm 4.7$ % | 12.6 / 63 | 10.0 / 50 | 38.0 / 190 | 35.4 / 177 |
| NuSVC (poly) | $79.4 \pm 7.5$ % | $75.2 \pm 5.0$ % | 12.2 / 61 | 10.6 / 53 | 37.4 / 187 | 35.8 / 179 |
| SVM (poly) | $79.2 \pm 10.1$ % | $75.7 \pm 7.0$ % | 11.6 / 58 | 11.2 / 56 | 36.8 / 184 | 36.4 / 182 |
| NuSVC (linear) | $70.5 \pm 8.1$ % | $64.9 \pm 5.9$ % | 19.0 / 95 | 14.6 / 73 | 33.4 / 167 | 29.0 / 145 |
| SVM (linear) | $68.9 \pm 9.5$ % | $66.6 \pm 3.9$ % | 16.8 / 84 | 15.6 / 78 | 32.4 / 162 | 31.2 / 156 |
| ***Ensemble methods*** | | | | | | |
| XGBoost | $75.8 \pm 5.6$ % | $76.0 \pm 3.3$ % | 11.0 / 55 | 11.8 / 59 | 36.2 / 181 | 37.0 / 185 |
| RandomForest | $74.5 \pm 5.1$ % | $78.2 \pm 5.2$ % | 9.6 / 48 | 12.4 / 62 | 35.6 / 178 | 38.4 / 192 |
| ***Probabilistic models*** | | | | | | |
| Naive Bayes (Gaussian) | $76.1 \pm 3.3$ % | $71.0 \pm 5.8$ % | 14.2 / 71 | 11.8 / 59 | 36.2 / 181 | 33.8 / 169 |
| ***Linear methods*** | | | | | | |
| Ridge Classifier | $74.4 \pm 8.1$ % | $66.3 \pm 5.6$ % | 18.8 / 94 | 12.8 / 64 | 35.2 / 176 | 29.2 / 146 |
| Logistic Regression | $73.3 \pm 9.0$ % | $65.5 \pm 5.7$ % | 18.6 / 93 | 13.6 / 68 | 34.4 / 172 | 29.4 / 147 |
| ***Baseline methods*** | | | | | | |
| K-Means (unsupervised) | $65.3 \pm 12.7$ % | $55.1 \pm 2.9$ % | 25.6 / 128 | 15.8 / 79 | 32.2 / 161 | 22.4 / 112 |
| Lin. Regression (thresholded) | $58.7 \pm 3.1$ % | $53.8 \pm 2.6$ % | 23.4 / 117 | 20.2 / 101 | 27.8 / 139 | 24.6 / 123 |

* “avg” refers to the per-fold average; “total” refers to the sum across all 5 folds. *Note: The results presented in this table are from supervised classifiers using balanced class weights.*

among the linear and nonlinear classifiers evaluated for KC classification, and with [3], who demonstrated that SVMs with non-linear kernels attain higher KC detection rates than linear kernel variants.

Table I summarizes the performance for each classifier tested in order of recall and precision. Average and Total number of False Positive (FP), False Negative (FN), True Positive (TP) and True Negative (TN) are also reported on this table.

### S2. CLASSIFICATION PERFORMANCE FOR EACH SUBJECT

The SVM classifier was evaluated on the Dataset 1 consensus-labeled dataset using 10-fold Group K-Fold cross-validation and the performance of the classifier across each subject was calculated. Table II presents the per-subject classification performance.

TABLE II
CLASSIFICATION PERFORMANCE FOR EACH SUBJECT IN DATASET 1: NUMBER OF TRUE POSITIVES (TP), FALSE POSITIVES (FP), FALSE NEGATIVES (FN), RECALL, SPECIFICITY AND ACCURACY FOR EACH SUBJECT AND IN TOTAL.

| Subject | N events | TP | FP | FN | Recall | Specificity | Accuracy |
|---|---|---|---|---|---|---|---|
| S23 | 38 | 14 | 2 | 5 | 73.7% | 89.5% | 81.6% |
| S24 | 44 | 21 | 8 | 1 | 95.5% | 63.6% | 79.5% |
| S25 | 62 | 23 | 7 | 8 | 74.2% | 77.4% | 75.8% |
| S26 | 58 | 24 | 7 | 5 | 82.8% | 75.9% | 79.3% |
| S27 | 36 | 10 | 7 | 8 | 55.6% | 61.1% | 58.3% |
| S28 | 86 | 37 | 10 | 6 | 86.0% | 76.7% | 81.4% |
| S30 | 38 | 17 | 5 | 2 | 89.5% | 73.7% | 81.5% |
| S31 | 32 | 15 | 3 | 1 | 93.8% | 81.3% | 87.5% |
| S32 | 46 | 21 | 6 | 2 | 91.3% | 73.9% | 82.6% |
| S33 | 40 | 14 | 3 | 6 | 70.0% | 85.0% | 77.5% |
| **Total** | **480** | **196** | **58** | **44** | **81.7%** | **75.8%** | **86.9%** |

Global metrics were computed from the aggregated confusion matrix: TP = 196, TN = 182, FP = 58 and FN = 44; and the classifier achieved a mean recall of 81.2% (± 12.0%; global of 81.7%) and specificity of 75.8% (± 8.2%; global of 75.8%), both intermediate between the two human experts, as discussed in the manuscript. Additionally, the classifier achieved the highest accuracy of 78.8% (± 7.4%; global of 86.9%), and F1-score of 79.3% (± 8.1%; global of 79.4%) across folds. As discussed on the manuscript, the majority of subjects achieved accuracy above 75%, with one notable exception: subject S27 (accuracy = 58.3%, recall = 55.6%, specificity = 61.1%).

### S3. EXTERNAL EVALUATION: UNION CRITERION OF DATASET 2

The SVM classifier trained on all 480 consensus-labeled samples from the Dataset 1 was applied to the Dataset 2 under the union criterion (5 excerpts, 210 KC, $2,398$ non-KC detected by localization algorithm). The model achieved a recall of $54.8\%$ and specificity of $88.1\%$ under the union criterion. In addition, to investigate the nature of classification errors, a morphological analysis applied to the primary dataset was performed. Figure 1 presents the average waveforms of false positives and false negatives alongside Dataset 2 and Dataset 1, KC and non-KC averages under the union criterion.

### S4. INTERNAL EVALUATION: UNION CRITERION OF DATASET 2

When the full pipeline was trained and evaluated within Dataset 2 using 5-fold Group K-Fold cross-validation with undersampling, performance improved substantially. Under the union criterion, global recall reached 71.0%, specificity 89.5%, and F1-score 48.8%. Figure 2 presents the average waveforms of false positives and false negatives alongside Dataset 2 true KC and true non-KC averages using the union labeling criterion.

Both external (Figure 1) and internal (Figure 2) evaluations supports what is mentioned in Section IV of the main manuscript: the classifier systematically detects events with genuine KC morphological characteristics that fall below the detection threshold of visual scoring. These both results also evidence that there are morphological differences in the candidate pools between Dataset 1 and Dataset 2.

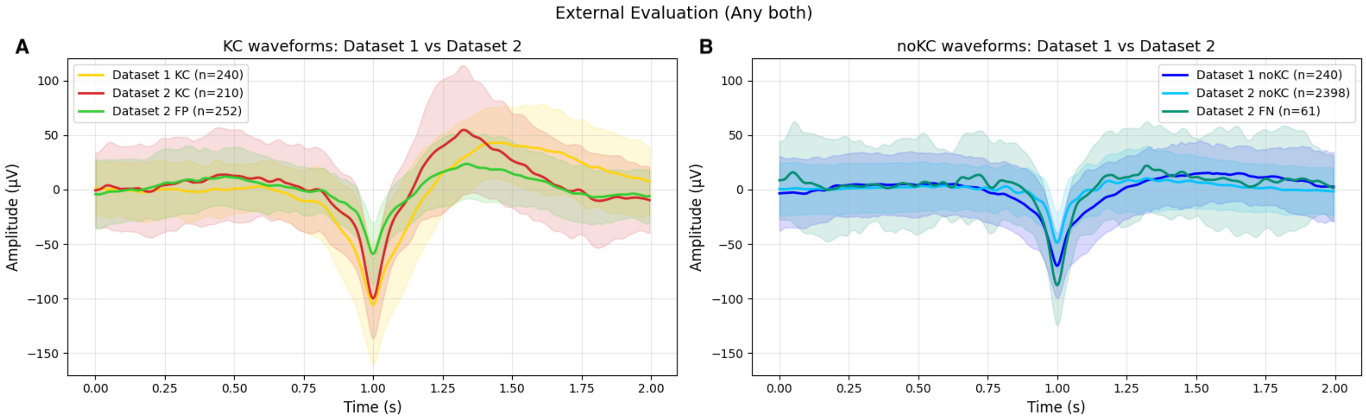


Fig. 1. External Evaluation on waveform morphology (Union criterion). **A** FP average compared against Dataset 1 and Dataset 2 KC averages. **B** FN average compared against Dataset 1 non-KC and Dataset 2 non-KC averages. Shaded regions represent $\pm 1$ SD.

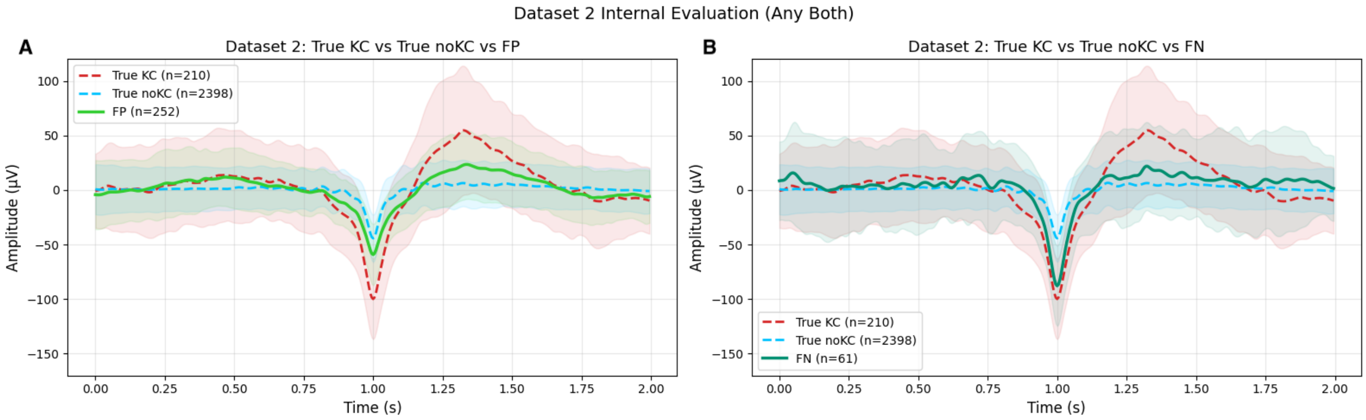


Fig. 2. Internal Evaluation on waveform morphology (Union criterion). **A** FP average compared against true KC and true non-KC averages. **B** FN average compared against true KC and true non-KC averages. Shaded regions represent $\pm 1$ SD.
